# Inferring Cancer Progression from Single-cell Sequencing while Allowing Mutation Losses

**DOI:** 10.1101/268243

**Authors:** Simone Ciccolella, Mauricio Soto Gomez, Murray Patterson, Gianluca Della Vedova, Iman Hajirasouliha, Paola Bonizzoni

**Affiliations:** Department of Computer Science, Systems and Communication, Univ. Milano-Bicocca, Milan, Italy; Institute for Computational Biomedicine, Department of Physiology and Biophysics, Weill Cornell Medicine of Cornell University, NY, USA; Englander Institute for Precision Medicine, The Meyer Cancer Center, Weill Cornell Medicine, NY, USA

## Abstract

**Motivation:** In recent years, the well-known Infinite Sites Assumption (ISA) has been a fundamental feature of computational methods devised for reconstructing tumor phylogenies and inferring cancer progressions seen as an accumulation of mutations. However, recent studies (Kuipers *et al.*, 2017) leveraging Single-cell Sequencing (SCS) techniques have shown evidence of the widespread recurrence and, especially, loss of mutations in several tumor samples. Still, established methods that can infer phylogenies with mutation losses are however lacking.

**Results:** We present the `SASC` (Simulated Annealing Single-Cell inference) tool which is a new and robust approach based on simulated annealing for the inference of cancer progression from SCS data. More precisely, we introduce a simple extension of the model of evolution where mutations are only accumulated, by allowing also a limited amount of back mutations in the evolutionary history of the tumor: the Dollo-*k* model. We demonstrate that `SASC` achieves high levels of accuracy when tested on both simulated and real data sets and in comparison with some other available methods.

**Availability:** The Simulated Annealing Single-cell inference (`SASC`) tool is open source and available at https://github.com/sciccolella/sasc.

## 1 Introduction

Recent developments in targeted therapies for cancer treatment rely on the accurate inference of the clonal evolution and progression of the disease. As discussed in several recent studies (Morrissy and Garzia, 2016; Wang *et al.*, 2016), understanding the order of accumulation and the prevalence of somatic mutations during cancer progression can help better devise these treatment strategies.

Most of the available techniques for inferring cancer progression rely on data from next-generation bulk sequencing experiments, where only a proportion of observable mutations from a large amount of cells is obtained, without the distinction of the cells that carry them. In recent years, many computational approaches have been developed for the analysis of bulk sequencing data with the purpose of inferring tumoral subclonal decomposition and reconstructing tumor phylogenies (evolutionary trees) (Strino *et al.*, 2013; Jiao *et al.*, 2014; Hajirasouliha *et al.*, 2014; Yuan *et al.*, 2015; Popic *et al.*, 2015; Malikic *et al.*, 2015; El-Kebir *et al.*, 2016; Marass *et al.*, 2016; Satas and Raphael, 2017; Bonizzoni *et al.*, 2017).

Single-cell Sequencing (SCS) technologies promise to deliver the best resolution for understanding the underlying causes of cancer progression. However, it is still difficult and expensive to perform SCS experiments with a high degree of confidence or robustness. The techniques available nowadays are producing datasets which contain a high amount of noise in the form of false negatives from allelic dropout, and missing values due to low coverage. Although this sequencing technology is rapidly improving, and some issues such as the presence of doublets are slowly fading away, it is important to develop methods that are able to infer cancer progression despite the lack of accuracy in the data produced by current SCS techniques.

Various methods have been recently developed for this purpose (Jahn *et al.*, 2016; Ross and Markowetz, 2016; Zafar *et al.*, 2017), some of them introducing a hybrid approach of combining both SCS and VAF (bulk sequencing) data (Ramazzotti *et al.*, 2017; Malikic *et al.*, 2017; Salehi *et al.*, 2017). Most of these methods, however, rely on the Infinite Sites Assumption (ISA), which essentially states that each mutation is acquired at most once in the phylogeny and is never lost. One reason for this is that such a simplifying assumption leads to a computationally tractable model of evolution, namely, the problem of finding a perfect phylogeny (Gusfield, 1991). This model is safe to use in settings such as the evolution of natural populations, and tends to be the norm more than the exception in this setting (Kimura, 1969). Cancer progression, however, is a fairly extreme situation, where the evolution is very fast, aggressive and with a high mutation rate. As a result, studies of SCS data are beginning to reveal phenomena that cannot always be explained with a perfect phylogeny (Kuipers *et al.*, 2017; Brown *et al.*, 2017). In (Kuipers *et al.*, 2017), the authors reveal widespread recurrence and loss of mutation, while in (Brown *et al.*, 2017), they find that large deletions on several branches of a tree can span a shared locus, and thus a given mutation may be deleted independently multiple times.

In this work we propose a novel and more general model to explain the above phenomena, which is not unnecessarily held back by strict adherence to the ISA. Some recent methods are beginning to appear, which have the same objective in mind, such as TRaIT (Ramazzotti *et al.*, 2017) and SiFit (Zafar *et al.*, 2017): in particular they allow deletions of mutations without specifying a particular model of evolution. In our approach, we use the Dollo model (Rogozin *et al.*, 2006), one of the models that is only slightly more general than the perfect phylogeny model, to allow the loss of point mutations. In particular, while it still constrains that a mutation can only be *acquired* at most once, it allows any number of independent losses of the mutation. Of course, the more general Dollo model departs from the convenient computational tractability of the perfect phylogeny model (Gusfield, 1991). However, if we restrict the number of losses of any mutation to 1 or 2 (rather than strictly 0), the resulting solution space is still small enough to explore a sizable portion of it in a reasonable amount of time, in practice.

Here we introduce the Simulated Annealing Single-cell inference (`SASC`) tool, a maximum likelihood phylogeny search framework that allows deletion of mutations, by incorporating the Dollo parsimony model (Farris, 1977). We show that our approach is competitive with the state-of-the art tools for inferring cancer progression from SCS data, while being the only tool to correctly identify important driver mutations in some real datasets, as verified by the manually curated progression scenarios for this data. In the next section, we layout the methodological basis of our tool, followed by Results and Discussion.

## 2 Methods

### 2.1 Formulation of the tree reconstruction problem

As mentioned before, cancer progression reconstruction can be modeled as the construction of a character-based incomplete phylogeny on a set of (cancer) cells, where each character represents a mutation.

In this framework we consider the input as an *n × m* ternary matrix *I_ij_*, where an entry *I_ij_* = 0 indicates that the sequence of cell *i* does not have mutation *j*, *I_ij_* = 1 indicates the presence of mutation *j* in the sequence of cell *i*, and a ? indicates that there is not enough information on the presence/absence of mutation *j* in cell *i.* This uncertainty about the presence of a mutation in a cell is a consequence of insufficient coverage in the sequencing.

However, the uncertainty of some entries is not the only issue that results from the sequencing process. In fact, entries of the input matrix *I* can also contain false positives and false negatives. We assume that these errors occur independently and uniformly across all the (known) entries of *I.* Namely, if *E_ij_* denotes the final *n × m* output matrix, i.e., the binary matrix without errors and noise estimated by the algorithm, then *α* denotes the false negative rate and *β* denotes the false positive rate. Hence, for each entry of *E_ij_* the following holds:

- *P* (*I_ij_* = 0|*E_ij_* = 0) = 1 *− β*
- *P* (*I_ij_* = 1|*E_ij_* = 0) = *β*
- *P* (*I_ij_* = 1|*E_ij_* = 1) = 1 *− α*
- *P* (*I_ij_* = 0|*E_ij_* = 1) = *α*

We aim to find a matrix which maximizes the likelihood of the observed matrix *I* (Jahn *et al.*, 2016) under the probabilities of false positives/negative and missing entries. That is we look for a matrix *E* that maximizes the following objective function

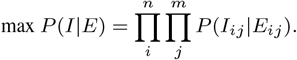

A more convenient but equivalent formulation can be obtained by taking the logarithm of the likelihood, obtaining then the following optimization problem:

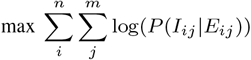

We point out that the values assigned to the unknown entries of the input matrix do not affect the objective function, that is *P* (*I_ij_* = ?|*E_ij_* =1) = *P* (*I_ij_* = ?|*E_ij_* = 0). To simplify the computation of the likelihood, we make a slight abuse and we pose *P* (*I_ij_* = ?|*E_ij_* = 1) = *P* (*I_ij_* = ?|*E_ij_* = 0) = 1.

However, since we are interested in the reconstruction of the evolutionary history of the input cells, then the resulting matrix *E* should contain clones that can be explained by an evolutionary process of the mutations. This restriction motivates the introduction of the phylogenetic tree concept.

A cancer phylogeny tree *T* on a set *C* of *m* mutations and *n* leaves is defined as a rooted tree whose nodes are labeled by the mutations of *C*, with the exception of the leaves, which are labeled as cells. Notice that the labeling must satisfy some restrictions depending on the evolutionary model that we consider. For example, in a perfect phylogeny tree no two nodes have the same label. This is a redefinition of classical character-based phylogeny, where the tree *T* is defined on a set of characters and where leaves have no label and represent different species.

The state of a node *x* is defined as the set of mutations that have been acquired but not lost in the path from the root to *x.* The state of each leaf *l* of *T* is naturally represented by an *m*-dimensional binary vector, called genotype profile, that we denote *D*(*T, l*), where *D*(*T, l*)_*j*_ = 1 if and only if the leaf *l* has the mutation *j* and zero otherwise.

We say that the tree *T encodes* a matrix *E* if there exists a mapping *σ* of the rows (cells) of *E* to the leaves of *T* such that for every row *i* of *E*, it follows that *E_i_* = *D*(*T, σ_i_*), where *σ_i_* denotes the image of row *i* by the mapping *σ.* Informally, *σ_i_* is the node in the phylogenetic tree corresponding to the parent of the cell *i.* Notice that the matrix *E* is fully characterized by the pair (*T, σ*).

In this manner, we can express the likelihood of the matrix *E* as: 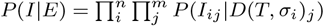, where the tree *T* encodes matrix *E.* Thus, our problem can be expressed as finding the solution of the following problem:

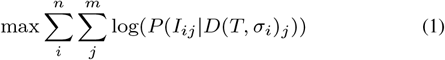

### 2.2 Introduction of the Dollo-*k* model

The Dollo parsimony rule assumes that in a phylogeny any single mutation is uniquely introduced in the evolutionary history but deletions of the mutation can occur any number of times.

The phylogeny reconstruction problem under a Dollo model is NP-complete (Day *et al.*, 1986). A restricted version of the Dollo model can be obtained by bounding the number of deletions for each mutation. We denote as Dollo-*k* the evolutionary model in which each mutation can be acquired exactly once and can be lost at most *k* times. The special cases Dollo-0 and Dollo-1 correspond to the perfect (Gusfield, 1991) and persistent (Bonizzoni *et al.*, 2016; Della Vedova *et al.*, 2017) phylogeny models, respectively. Since the Dollo evolutionary model allows back mutations, we introduce a new type of node label in the phylogenetic tree, to take into account the mutation losses. For each mutation *p* we create *k* new mutations 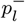for *l* ∈ {1, …, *k*}, representing the possible losses of mutation *p.* As in the perfect case, we require that no two different nodes have the same label. Additionally, we impose that all nodes labeled by a mutation loss *p*^−^ are descendants of the node labeled by mutation gain *p.* Consequently, the vector *D*(*T, σ_i_*) which expresses the genotype profile of a row *i* will have a one in mutations acquired but never lost in the path from the root to the parent *σ_i_* of the leaf *i.* We stress the fact that, unlike the case of the perfect phylogeny, when deletions are introduced the set of trees encoding a given solution could have more than one element. We can see that switching the labels of nodes *b*^−^ and *d*^−^ in Figure 1 produces a different tree which is still a solution of the proposed input matrix when the Dollo model is considered. Moreover, we see that the ancestral relationships between those two mutations is opposite in both representations. When the number of cells, mutations and possible deletions increase, and with the noise caused by false calls and missing entries, this problem is amplified, increasing the number of different cancer progression phylogenies which equally explain the same input. A more complex example can be seen in Figure 2 where a different order of mutations and a different set of deletions can equally explain a given input.

**Fig. 1.**
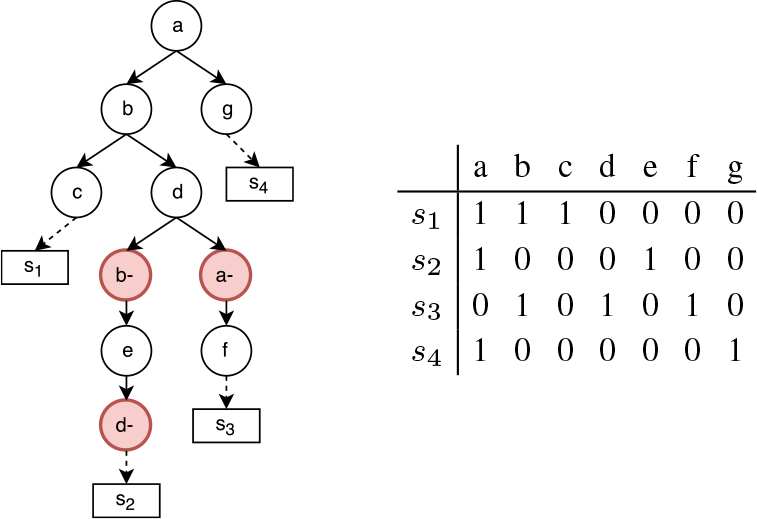
Example of a binary matrix that does not allow a perfect phylogeny, since columns *a* and *b* are in conflict, i.e. the four gametes rule (Gusfield, 1991) does not hold. The tree represents one of the possible Dollo phylogenies that explain the matrix.

**Fig. 2.**
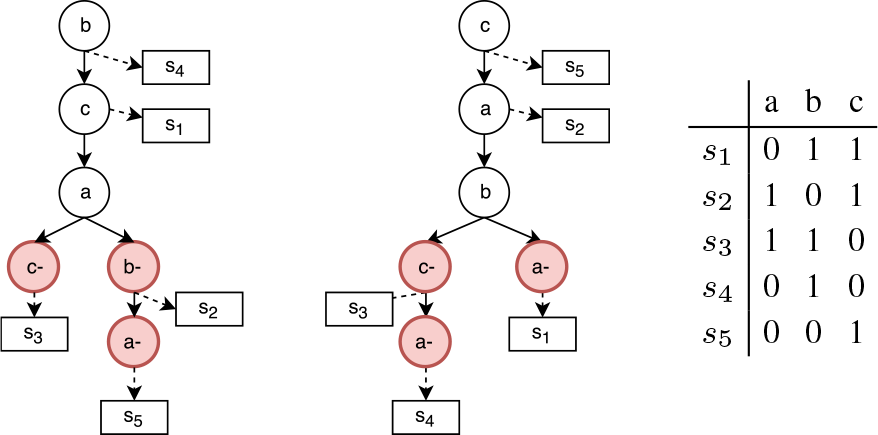
Example of two Dollo phylogenies that explain the same binary matrix. It is important to notice that the ancestral order of mutations *c, a* and *b* is inverted but the two different trees can equally explain the input binary matrix. In fact, in a Dollo phylogeny the order of two mutations can be inverted and, thank to the introduction of deletions, they could both be correct solutions for a given input.

### 2.3 Simulated Annealing

As mentioned before, the fact that (1) we can flip entries, and (2) we want to find the maximum likelihood tree, makes the phylogeny reconstruction problem under the Dollo-*k* model computationally hard for any *k* > 0. For this reason we consider the Simulated Annealing (Kirkpatrick *et al.*, 1983) (SA) approach in order to find a tree which maximizes the likelihood of an incomplete input matrix and that satisfies the Dollo-*k* phylogeny model, where *k* is given as input.

SA is a random search technique which explores the region of feasible solutions searching for an optimal one. Unlike other deterministic search methods which can be trapped into local optima, SA has the ability to overcome this drawback and converge to a global optimum. The basic idea of the algorithm is to perform a random search which accepts, with some probability, changes that not necessary improve the objective function. At each step, the probability of moving to some state with a smaller value changes according to a parameter called the *temperature*, which continuously decreases as the exploration evolves. In the first iterations of the algorithm execution, the temperature is very high, and it is possible (with a fairly large probability) to accept a move into a state with a lower objective value, but as temperature decreases, the probability of moving also decreases. At the end, when the temperature is sufficiently low, the algorithm becomes a local search method, hence unable to escape a local optimum.

#### 2.3.1 Neighborhood topology

When the feasible region topology is not explicitly defined, we must provide the way in which the algorithm search process can move from a given state to another. In our particular framework we attempt to find a tree, thus we must define the neighborhood of a phylogenetic tree in the feasible region. This choice is crucial in the algorithm definition since it determines the way in which feasible solutions are explored and ultimately determining whether or not the algorithm converges.

Before detailing the SA algorithm we describe the valid tree operations that define if two phylogenetic trees are neighbors in the search space which defines the set of possible simulated annealing moves for each iteration. For the sake of clarity we introduce some notation: given a phylogenetic tree *T* and a node (labeled as) *i*, the *ρ*(i) denotes the father of *i* in *T.*

- **Subtree Prune and Regraft (SPR) (Swofford and Olsen, 1990)**: given a tree *T* and two internal nodes *u, v* ∈ *T* such that neither is an ancestor of the other, we prune the subtree rooted in *v* by removing the edge (*v, ρ*(*v*)) and we grafted as a new child of *u* by adding the edge (*v, ρ*(*u*)).
- **Add a deletion**: given two nodes *u, v* ∈ *T* such that *v* is an ancestor of *u*, we insert a node *v*^−^ that represents a loss of mutation *v.* The new node is made the parent of *u.* We remark that this operation takes place only if the resulting tree satisfies the desired phylogeny model. More precisely, for the Dollo-*k* we must check that the mutation *v* has been previously lost in the tree at most *k −* 1 times, never lost in any ancestor or descendant of *v*^−^.
- **Remove a deletion**: given a node *u* ∈ *T*, labeled as a loss, we simply remove it from the tree *T*: all children of *u* are added as children of *ρ*(*u*) and the node *u* is then deleted.
- **Swap nodes labels**: given two internal nodes *u, v* ∈ *T* the labels of *u* and *v* are swapped. If a previously added loss becomes invalid due to this operation — because a mutation *c* is lost in a node *c*^−^, but the node where the mutation *c* is acquired is not an ancestor of *c*^−^ any more — then we removed the deletion *c*^−^.
- **Change cell assignment (CCA)**: unlike that previous operations which modify the shape of the phylogenetic tree, this operation works on the label assignment *σ.* Given a leaf *l* and a node *u*, randomly chosen, we redefine the father of *l* as *u.* That is, we remove the edge (*l, ρ*(*l*)) and we create the edge (*l, u*); in other terms we set *σ_l_* = *u.*

#### 2.3.2 The algorithm

The algorithm works in two phases. In the first phase, the goal of the algorithm is to find a maximum likelihood perfect phylogeny tree, while in the second phase, deletions are added in order to improve the solution by constructing a Dollo-*k* phylogeny. In each phase, a SA process is performed using a different set of valid operations but according to the the same temperature decay process.

In the **First Phase**, `SASC` searches for the maximum likelihood perfect phylogeny tree by modifying subtrees using SPR and CCA operations. In each iteration one of these operations is performed with the same probability.

In the **Second Phase** we extend the solution found in the first phase by adding possible losses without radically change the topology of the tree *T.* Thus, valid operations are add/remove deletions, swap node labels and CCA. In each iteration, one of this operations is performed with equal probability.

In both phases the nodes in which valid operations are performed are chosen uniformly among all possible candidates. Moreover, in both SA search processes we have that given a tree and a valid tree operation, the probability of accept the new solution the probability of acceptance as min{e^∆^*^v/T^*, 1}, where ∆*v* is the possible change in the likelihood function after performing the operation and *T* is the current temperature. The **cooling process** follows a geometric decay with a factor of 10^*−*5^, i.e.the temperature at the *i*-th iteration is equal to *T_i_* = (1 *–* 10^*−*5^)*T_i−_*_1_ and *T*_0_ = 10^5^. The SA process stops when the temperature drops below a lower bound set at 10^−4^. All previously mentioned parameters were empirically found to be the best for both accuracy and running time in our experimentation.

Since mutation losses are not as frequent as mutation gains, our approach allows to set an upper bound on *d*: the total number of deletions of the resulting tree. For example, in a Dollo-*k* model we can consider only trees where each mutation is lost at most *k* times, but there are at most *d* nodes associated to mutation losses.

## 3 Results

### 3.1 Results on simulated data

We have tested our method on simulated data, where the ground truth phylogeny tree is known. We recall that it is possible, however, that a completely different tree achieves a better likelihood on the input data than the one obtained via simulation. This problem is essentially unavoidable, since generating a progression that is the unique solution for the corresponding SCS input matrix would require the contrived addition of artifacts to both the desired tree and the input matrix, in order to guarantee this uniqueness. These artifacts would likely be so artificial that resulting instance would not satisfy even the basic assumptions on cancer progression.

#### 3.1.1 Generating simulated datasets

We have simulated two datasets, according to the parameters of Table 1. For each dataset we randomly generated 50 clonal trees. Given the number *S* of subclones, we generated a random tree of *S* nodes by adding a new node as a child of a random pre-existing one. Each of the *M* mutations *q*_1_, …, *q_M_* is then, uniformly at random, assigned to one of the *s_i_* subclones. We allowed at most a fixed number *d* of deletions in each clonal tree: therefore *d* new nodes are added to the tree at random positions. A deletion of a mutation is then assigned to each of the *d* new nodes, by picking uniformly at random, one of the mutations which is affecting the parent of the node and which has not been already lost in the path, according to the Dollo model.

**Table 1.** Parameters used to simulate the input matrices, where *S* is the number of subclones, *M* is the number of mutations, *N* is the number of cells, *d* is the maximum number of allowed mutation deletions, *α* is the false negative ratio, *β* the false positive ratio and *γ* is the missing data ratio.

| Experiment | $S$ | $M$ | $N$ | $d$ | $\alpha$ | $\beta$ | $\gamma$ |
| --- | --- | --- | --- | --- | --- | --- | --- |
| 1 | 7 | 30 | 150 | 3 | 0.15 | $10^{-3}$ | 0.25 |
| 2 | 9 | 30 | 100 | 5 | 0.1 | $10^{-4}$ | 0.1 |

To obtain the genotype profile of the *N* cells, we uniformly assigned at random each cell to a node and derived its profile from the clonal tree. Finally to simulate noise in the data, we flipped a 0 entry to 1 with probability *β* to simulate false positives and a 1 entry to 0 with probability *α* to simulate false negatives. Moreover, each entry has a probability *γ* to be a missing entry. All flips and missing values are uniformly and independently distributed, without repetitions.

#### 3.1.2 Evaluation on simulated datasets

We measured the accuracy of `SASC` with two scores based on standard cancer progression measures used in various studies (Malikic *et al.*, 2017; Jahn *et al.*, 2016) and a novel parsimony based score, defined as follows:

**Ancestor-Descendant accuracy**: This measure considers all pairs of mutations (*x, y*) that are in an Ancestor-Descendant relationship in the ground truth tree *T.* For each such pair we check whether the ancestor-descendant relationship is conserved in the inferred tree *I*, in fact we calculate the number of mutations in a Ancestor-Descendant correctly inferred (true positives); the number of mutations that are incorrectly inferred to have an Ancestor-Descendant relationship (false positives); the number of mutations correctly inferred to not be Ancestor-Descendant (true negatives); finally the number of mutations incorrectly inferred to not have an Ancestor-Descendant relationship (false negatives). The score is defined by the *F*_1_ score, that is the geometric mean of the precision and recall.

**Different-Lineage accuracy**: Just as the previous measure, we consider all pairs of mutations (*x, y*) that are not in an ancestor-descendant relationship, i.e. are in different branches of *T.* The score is defined, similarly to the previous measure, as the resulting *F*_1_ score.

**Parsimony score**: This is a parsimony-based measure. We measure the difference between the number of flips, i.e., changes from 0 to 1 and from 1 to 0, estimated by some tool to correct the input, and the actual number of flips introduced by the simulation process to induce the noise. The rationale is that a good solution should be smaller, i.e., closer to the correct amount of changes introduced by the simulation process. Formally, the parsimony score is defined as |*H* (*S*) *− H* (*E*)| where *H* (*S*) is the total number of flips induced by the simulation, and *H* (*E*) is the number of flips estimated by the tool. While this measure does not consider the overall accuracy of a solution, it is a good estimation if used in conjunction with the previous ones.

Note that none of the above mentioned metrics accounts for ISA violations. We decided to compare `SASC` against SCITE (Jahn *et al.*, 2016) and SiFit (Zafar *et al.*, 2017): while B-SCITE (Malikic *et al.*, 2017) is a clear improvement over SCITE, it combines single-cell data with bulk sequencing data — since we do not manage the latter kind of data, a fair comparison is not feasible. For the same reason we do not compare against TRaIT (Ramazzotti *et al.*, 2017). OncoNEM (Ross and Markowetz, 2016) was excluded because it infers cell lineage progressions instead of mutational progressions, therefore it is not possible to compare our predictions with theirs. Furthermore, OncoNEM does not complete the execution on datasets as large as the ones used in the simulations.

Figure 3 shows the comparison between `SASC`, SCITE, and SiFit for *Experiment 1.* The performance obtained by `SASC` for the *Ancestor-Descendant* and the *Different Lineages* measures is essentially the same as SCITE, while both tools obtain better trees than SiFit. `SASC` shows a high accuracy (94%) on the detection of deletions. Both SCITE and SiFit do not detect any deletion but only true negatives. However, while SCITE’s outcome was expected, since it was not designed to detect ISA violations, SiFit was not able to detect any violation in any of our simulations, despite having been developed for this scope. Lastly, Figure 4 shows the *Parsimony Score* where we can see `SASC` and SCITE scoring very similarly, while SiFit requires a larger number of flips.

**Fig. 3.**
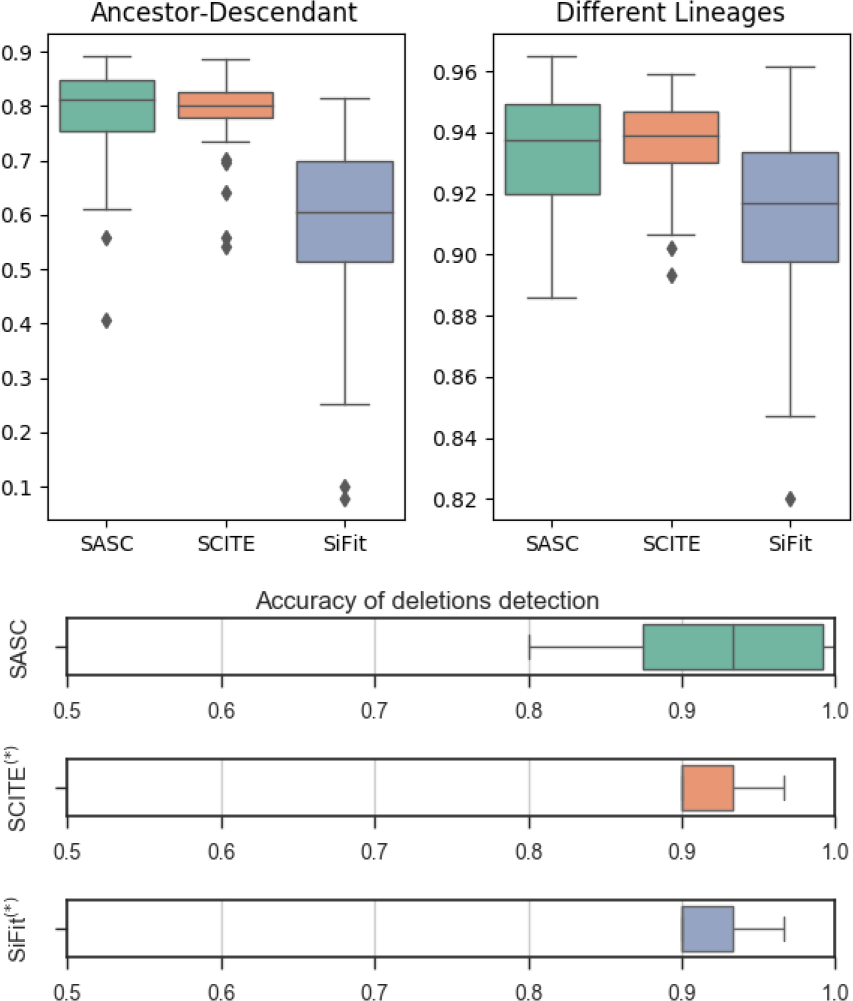
Accuracy results for Experiment 1, described in Section 3.1.2. `SASC` and SCITE are relatively close in the Ancestor-Descendant and Different Lineages measures, with `SASC` achieving a slightly better accuracy for the former and the opposite for the latter. On the other hand, SiFit achieves the poorest result for the Ancestor-Descendant measure and the best for the Different Lineages, this is possibly to due to the tendency of branching in the model. The lower plot shows the standard accuracy in deletion calling, i.e.the number of mutation correctly classified as deletion and not deleted over the total population for all the tools; (*) SCITE and SiFit detect no deletion, therefore they are correctly classifying only true negatives and thus they achieve the same score.

**Fig. 4.**
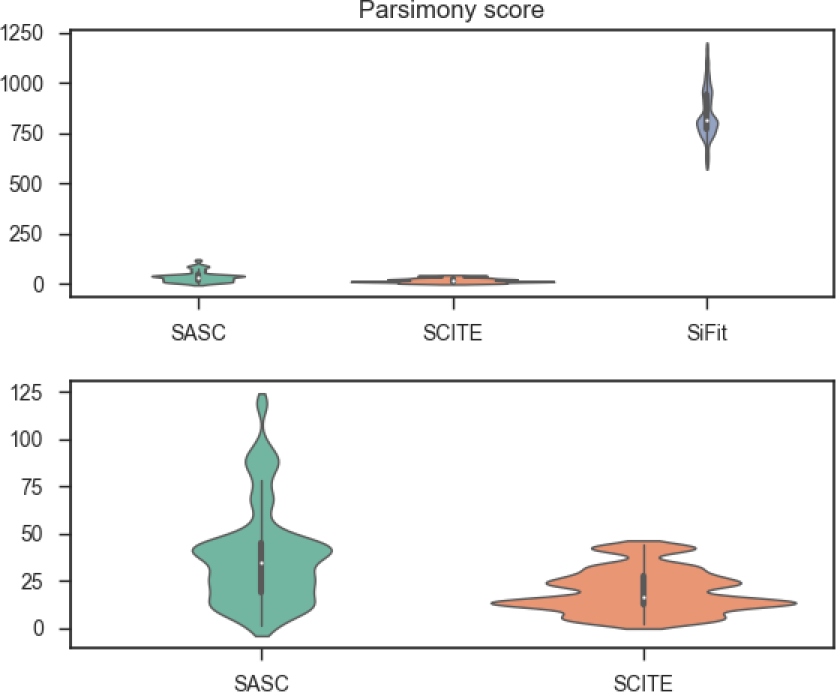
Parsimony scores for the input matrix for Experiment 1. The upper plot shows the parsimony scores for `SASC`, SCITE and SiFit, calculated as the absolute value of the difference between the total number of flips (from 1 to 0 or vice versa) introduced in the simulations and the total number of flips introduced by the corresponding tool. See Section 3.1.2 for a detailed definition of parsimony score. The lower plot is focused on the results for `SASC` and SCITE. The tools achieve a similar accuracy, with `SASC` being consistently closer to the correct number of flips while presenting a few more outliers than SCITE.

Results for *Experiment 2*, shown in Figures 5 and 6, corroborate the results of the previous experiment: `SASC`, SCITE and SiFit score similarly on the tree-based measure. In this experiment SCITE gives slightly better results than `SASC`, while both tools outperform SiFit. Again, `SASC` is the only method capable of detecting deletions, with a 93% accuracy. Lastly `SASC` and SCITE have similar values of *Parsimony Score*, SiFit has higher values.

**Fig. 5.**
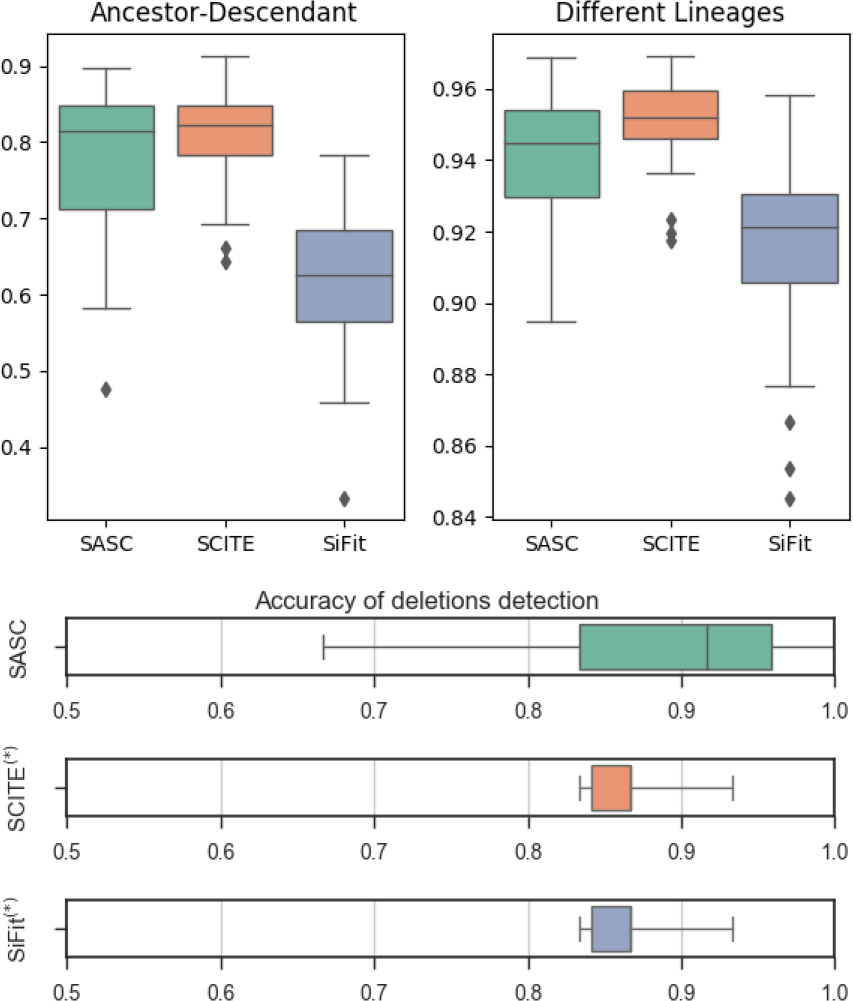
Accuracy results for Experiment 2, described in Section 3.1.2. `SASC` and SCITE are relatively close in the Ancestor-Descendant and Different Lineages measures, with `SASC` achieving a slightly better accuracy for the former and the opposite for the latter. On the other hand, SiFit achieves the poorest result for the Ancestor-Descendant measure and the best for the Different Lineages, this is possibly to due to the tendency of branching in the model. The lower plot show the standard accuracy in deletion calling, i.e.the number of mutation correctly classified as deletion and not deleted over the total population for all the tools; (*) SCITE and SiFit detect no deletion, therefore they are correctly classifying only true negatives and thus they achieve the same score.

**Fig. 6.**
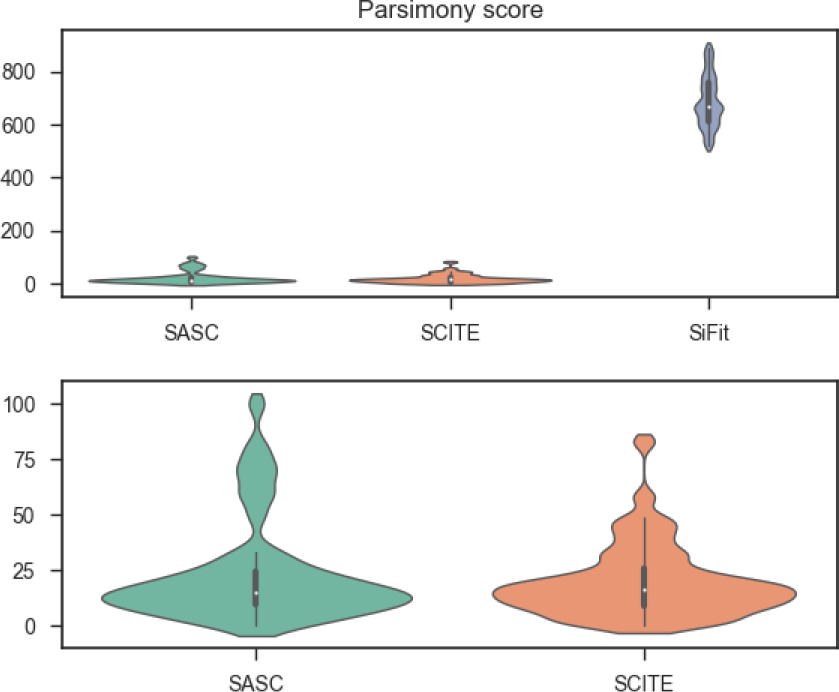
Parsimony scores for the input matrix for Experiment 2. The upper plot shows the parsimony scores of `SASC`, SCITE and SiFit, calculated as the absolute value of the difference between the total number of flips (from 1 to 0 or vice versa) introduced in the simulations and the total number of flips introduced by the tool. See Section 3.1.2 for a detailed definition of parsimony score. The lower plot is focused on the results for `SASC` and SCITE. The tools achieve a very similar accuracy, with `SASC` presenting slightly more outliers.

### 3.2 Results on real cancer data

We tested `SASC` on Childhood Acute Lymphoblastic Leukemia data from (Gawad *et al.*, 2014). In particular, we focused on Patient 4 and Patient 5 of this study, given their large amount of both cells and mutations, as well as their complexity. Data on Patient 4 consists of 78 somatic Single Nucelotide Variants (SNVs) over 143 cells, while Patient 5 is affected by 104 somatic SNVs over 96 cells. The original study estimated an allelic drop-out rate of less than 30%. Since the trees in (Gawad *et al.*, 2014), determined using expectation maximization on a multivariate Bernoulli distribution model, are manually curated and of high quality, we select them as the ground truth.

To ensure the absence of doublets, i.e. noise produced by error due to the fact that two cells are sequenced instead of single-cell, we preprocessed the input using the *Single-cell Genotyper* (SCG) tool (Roth *et al.*, 2016). SCG is a statistical model which removes all cells of the datasets that are likely to be doublets. Since SCG is reliable, we focus on doublet-free data in the design and experimental analysis of `SASC`. Moreover, doublets are becoming rarer as single-cell technology progresses.

Figure 7 shows the tree inferred by `SASC` for Patient 4; `SASC` correctly infers the tree structure assumed in the study as well as the size of subclonal the population. The driver mutations are correctly detected, and mutations COL5A2, SDPR and TRHR are inferred as deletions. Furthermore boldfaced and colored mutations indicate the correctly inferred specific driver mutations for the subclone of the same color. It is interesting to notice that the violet clone is supposed not to have mutation COL5A2: this particular mutation is indeed deleted in the clone. This solution was found assuming a Dollo-2 phylogeny model with no restriction on the total number of deletions in the cancer progression. Figure 8 shows the tree inferred by SCITE for the same dataset, the tree structure is similar to the one proposed in the paper but it presents more clones. Furthermore we highlighted in red driver mutations that were not correctly detected and in blue mutations that define a subclone and should be in the same cluster.

**Fig. 7.**
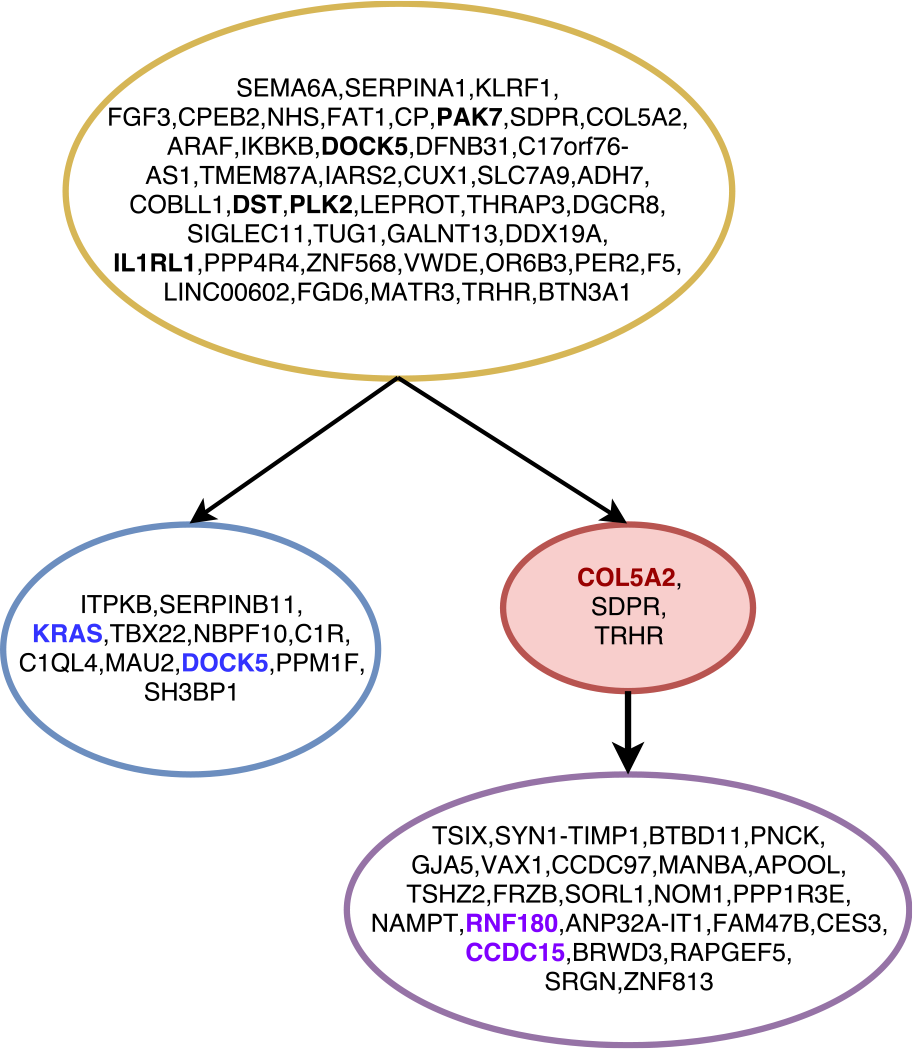
The tree inferred by `SASC` for Patient 4 of Childhood Lymphoblastic Leukemia data from (Gawad et al., 2014). Different clones are indicated with different colors. Red nodes indicate deletions of mutations, while boldfaced mutations are the mutations indicated as driver in the original sequencing study. Mutations in bold and colored are driver mutations for the clone with the same color. Mutations are clustered by collapsing simple linear paths.

**Fig. 8.**
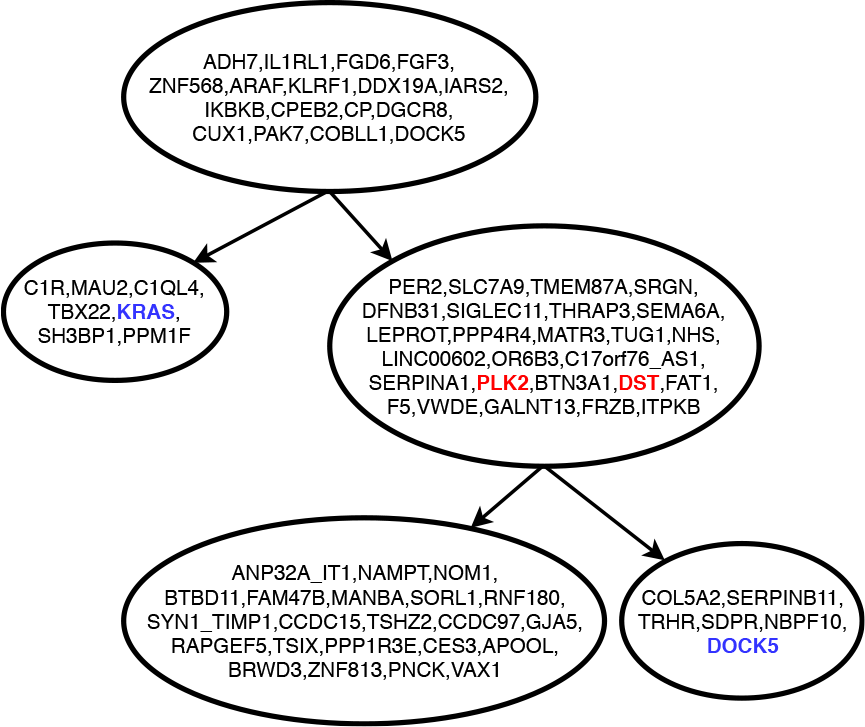
The tree inferred by SCITE for Patient 4 of Childhood Lymphoblastic Leukemia data from (Gawad et al., 2014). Mutations highlighted in red are driver mutations not correctly detected, while mutations highlighted in blue are two mutations that define a subclone and should be in the same cluster. Mutations are clustered by collapsing simple linear paths.

In Figure 9, the inferred solution for Patient 5 of the same study is shown. As in the previous dataset, our inferred tree perfectly supports the hypotheses proposed in the sequencing study: in fact, it correctly infers the topology of the tree, as well as the driver mutations. Boldfaced mutations are the driver mutations for the tree or the subclone with the same color. This solution was found assuming a Dollo-2 phylogeny model with a restriction of 10 deletions in the cancer progression, as described in Section 2.3. Figure 10 shows the tree inferred by SCITE for Patient 5 of the study; the tree topology is correctly inferred, however mutations highlighted in red are driver mutations that were not correctly detected.

**Fig. 9.**
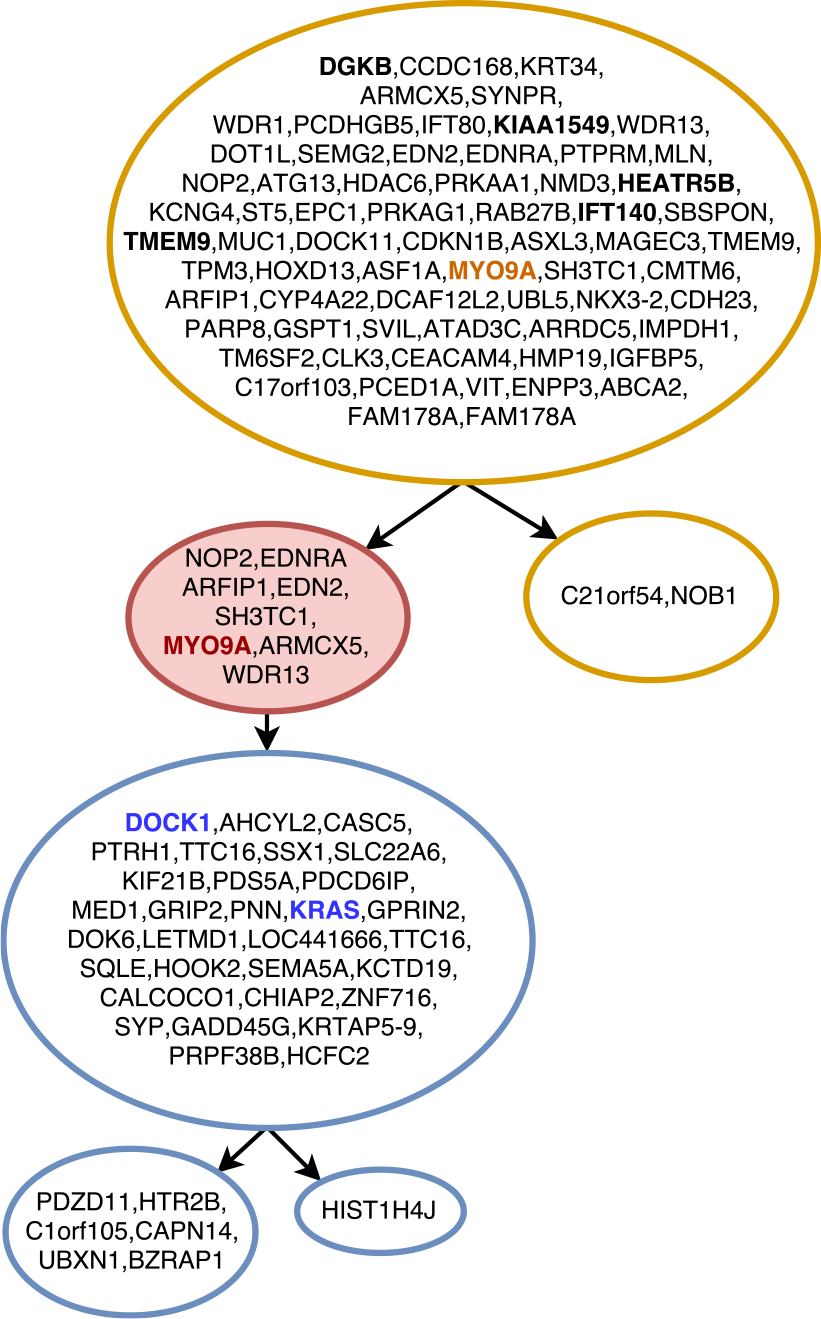
Tree inferred by `SASC` for Patient 5 of Childhood Lymphoblastic Leukemia data from (Gawad et al., 2014). Different clones are indicated with different colors while the red-colored nodes indicate deletions of mutations and mutations highlighted in bold are the mutations indicated as driver in the original sequencing study. Mutations boldfaced and colored are driver mutations for the same colored clone. Mutations are clustered by collapsing simple linear paths.

**Fig. 10.**
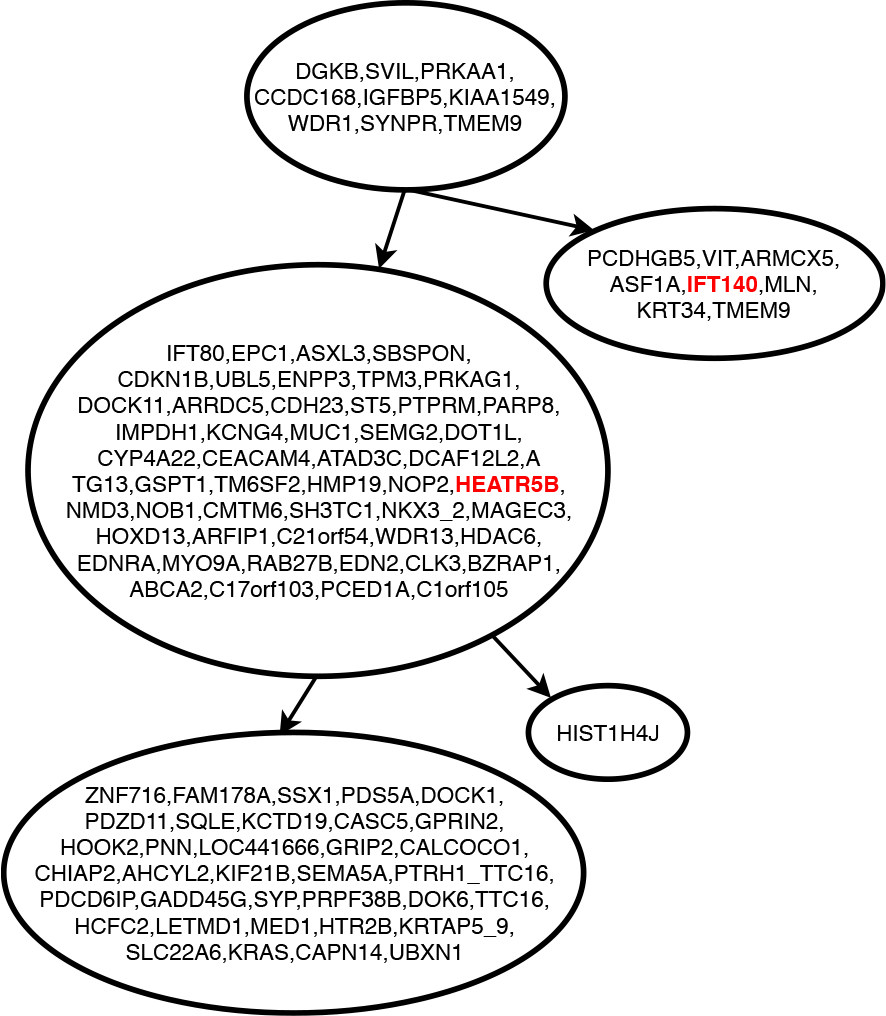
Tree inferred by SCITE for Patient 5 of Childhood Lymphoblastic Leukemia data from (Gawad et al., 2014). Mutations highlighted in red are driver mutations not correctly detected. Mutations are clustered by collapsing simple linear paths.

We also tested `SASC` on a recent Single-cell RNA-seq sequencing study of primary Breast Cancer (Chung *et al.*, 2017). Figure 11 represents the inferred cancer progression of Patient 9 of the study. For each mutation studied in (Chung *et al.*, 2017), its Raw Transcripts Per Kilobase Million (TPM) value is publicly available on the NCBI Gene Expression Omnibus database. We selected the mutations whose TPM is at least 10000: this resulted in 42 mutations. Moreover we have considered only the 60 cells of Patient 9 that have at least one of the selected mutations. The sequencing study (Chung *et al.*, 2017) does not propose a clonal tree, however several deletions were expected, since it is typical of genetic alterations in breast cancer. We ran `SASC` with two different configurations: first we did not limit the number of mutations under a Dollo-2 model and it inferred a total of 20 deletions. Then, for a clearer visualization (represented in Figure 11) we allowed only 5 deletions. Furthermore B2M, which is considered a driver mutation (Rajendran and Deng, 2017), is correctly detected as the chronologically first mutation. SCITE infer a tree, shown in Figure 12 very similar to the one inferred by `SASC` and both tools detect the same set of driver mutations.

**Fig. 11.**
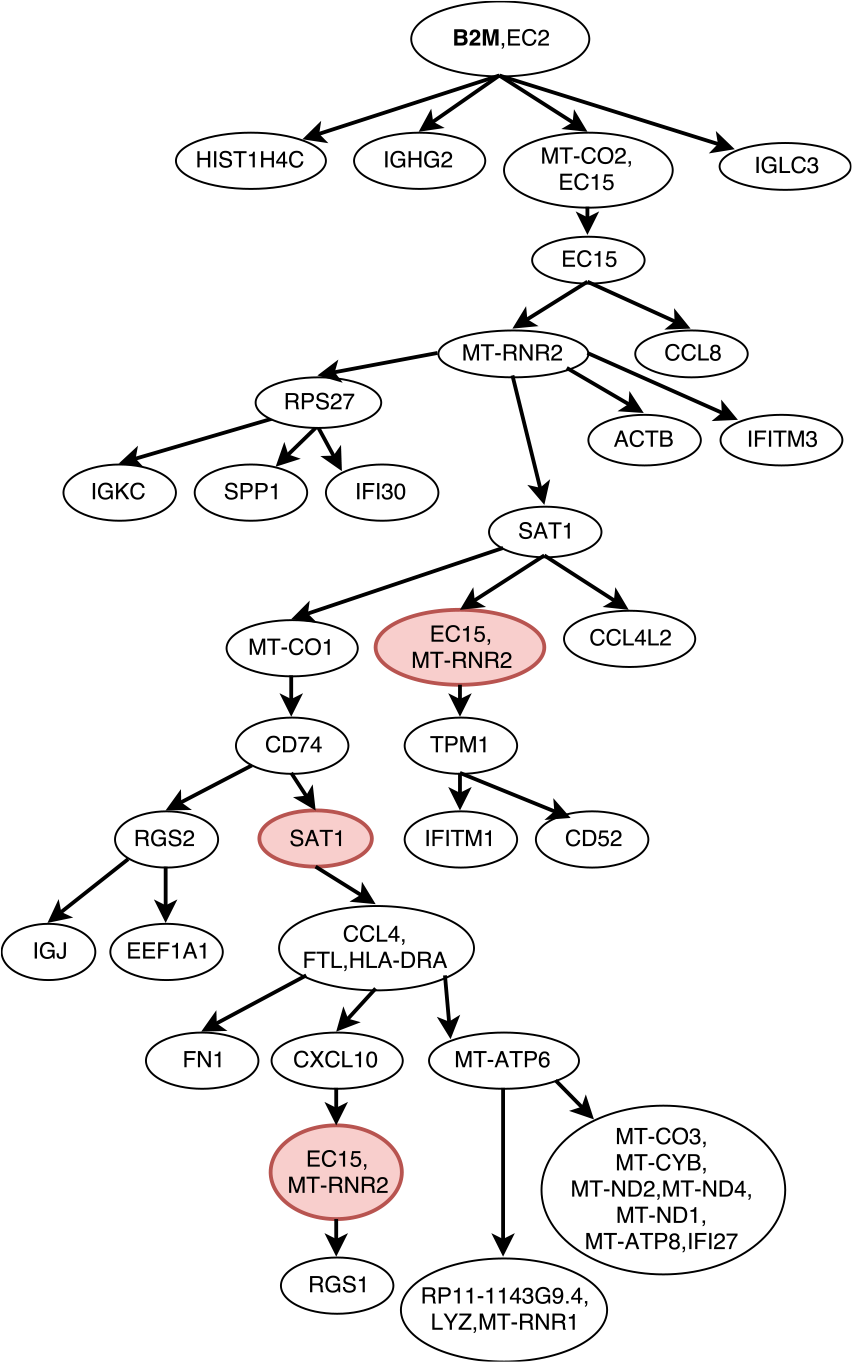
Tree inferred by `SASC` for Patient 9 of primary Breast Cancer data from (Chung et al., 2017). Red nodes indicate a deletion of mutations, while mutations highlighted in bold are the mutations indicated as driver.

**Fig. 12.**
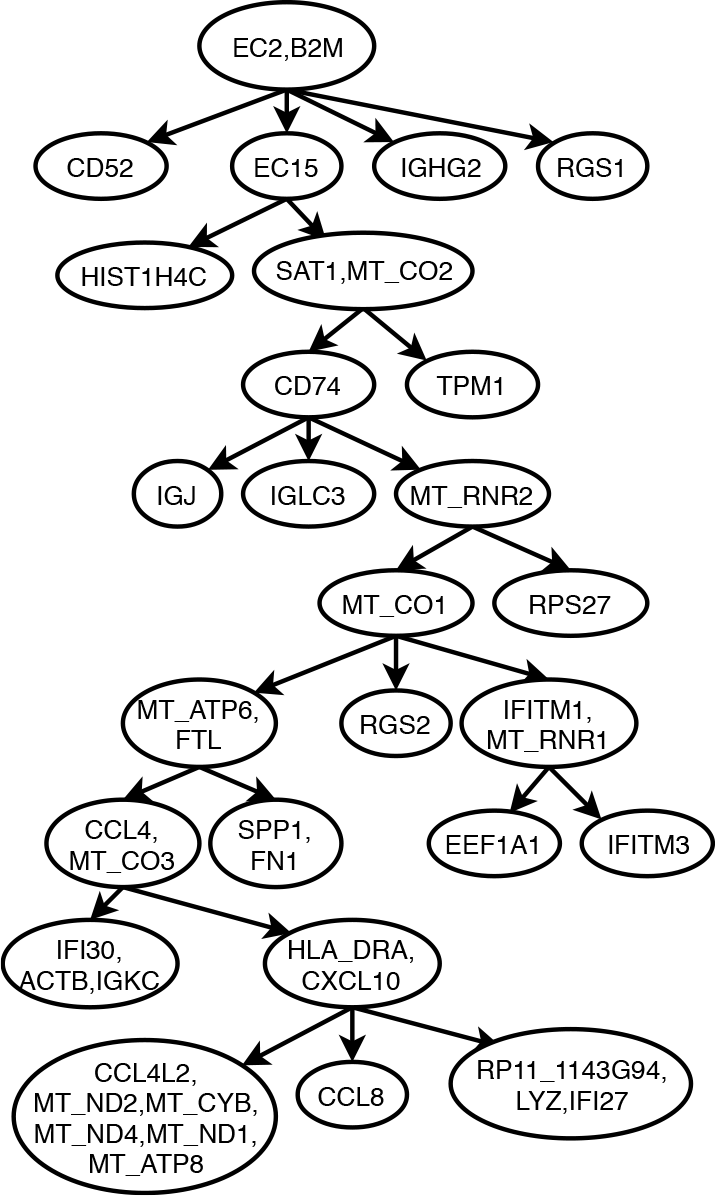
Tree inferred by SCITE for Patient 9 of primary Breast Cancer data from (Chung et al., 2017). The tree structure is very similar to the one inferred by `SASC`, and driver mutations are correctly detected by both tools.

## 4 Discussion

`SASC` is an accurate tool for inferring intra-tumor progression and subclonal composition from SCS data, and it is robust to various sources of noise in these data. `SASC` is highly accurate on simulated data where it scores better than, or very similarly to, SCITE.

On real data, `SASC` performs extremely well and it infers correctly the expected phylogeny tree structure, as well as the driver mutations and the decomposition of the clones. Furthermore, it can be used on very large datasets. Since the actual value of the parameters *α* and *β* are unknown, we suggest to try different input values for the parameters *α* and *β*: they affect the overall solution and can lead to different sets of solutions. A particularly interesting example is given by the inferred tree in Figure 9. The corresponding input dataset in this case contains more than 5000 conflicts between mutations – according to the four gametes rule, each one witnessing a violation of the ISA, by definition. With only a slight relaxation of the ISA — the Dollo-2 model — `SASC` is able to infer an accurate solution with a total of only 10 deletions, while perfect phylogeny methods would require a large amount of flips of the entries just to produce a feasible solution in this case.

As a consequence of its two-phase algorithm, `SASC` first finds the maximum likelihood perfect phylogeny solution and only later, in the second phase, does it augment it to a Dollo phylogeny. Therefore if it is not possible to introduce a deletion that improves the likelihood, then the method will produce a perfect phylogeny. Hence, `SASC` only employs the more general Dollo model only when it can improve the overall solution.

In summary, by way of its accuracy, `SASC` provides new insights into the analysis of intra-tumor heterogeneity by proposing a new progression model that has never been previously applied in cancer phylogeny reconstruction on Single-cell data.

## Acknowledgements

SC acknowledges the support of a Mobility Exchange Fellowship from the University of Milano-Bicocca. Part of this work has been done during a visit by SC to Weill Cornell Medicine.

